# Renal Interstitial Cells Promote Nephron Regeneration by Secreting Prostaglandin E2

**DOI:** 10.1101/2021.08.27.457936

**Authors:** Xiaoliang Liu, Ting Yu, Xiaoqin Tan, Daqing Jin, Jiangping Zhang, Yunfeng Zhang, Shuyi Liao, Wenmin Yang, Jinghong Zhao, Tao P Zhong, Chi Liu

**Author notes:** X. L., and T.Y. contributed equally to this work.

## Abstract

In organ regeneration, progrnitor and stem cells reside in their native microenvironment, which provides dynamic physical and chemical cues essential to their survival, proliferation and differentiation. However, what kind of cells provide a native microenvironment for renal progenitor cells has not been clarified. Here, single-cell sequencing of zebrafish kidney revealed that *fabp10* was a marker of renal interstitial cells (RICs), and the *Tg(fabp10a:GFP)* transgenic line can specifically label RICs in zebrafish kidney. The formation of RICs and nephrons are closely accompanied during nephron regeneration. RICs form a network to wrap the renal progenitor cell aggregates. RICs in close contact with cell aggregates express cyclooxygenase 2 and secrete prostaglandin 2 (PGE2). Inhibiting PGE2 production prevented nephrogenesis by reducing the proliferation and differentiation of progenitor cell aggregates. PGE2 promoted maturation of the nephron by activating the WNT signaling pathway in progenitor cell aggregates in cooperation with Wnt4a. These findings suggest that RICs provide a necessary microenvironment for rapid nephrogenesis during nephron regeneration.

## Introduction

Acute renal injury (AKI) is the most common cause of organ dysfunction in critically ill adults, and the in-hospital mortality is as high as 62% (Doyle & Forni, 2016). However, the mechanisms of how kidney recover rapidly from injury have not been elucidated. Unlike mammals, zebrafish can regenerate a large number of nephrons rapidly after kidney injury to make up for lost renal function (Diep et al., 2011). During regeneration, *lhx1a*-positive (*lhx1a*^+^) progenitor cells congregate to form cell aggregates, then proliferate and differentiate into renal vesicles, which finally develop into mature nephrons (Diep et al., 2011). Progenitor and stem cells reside in their native microenvironment, which provides dynamic physical and chemical cues essential to their survival, proliferation and function (Lee, Hirasawa, Garcia, Ramanathan, & Kim, 2019). However, what kind of cells provide a native microenvironment for renal progenitor cells has not been clarified.

RICs are specialized fibroblast-like cells found mainly between the tubular and vascular structures (Whiting, Tisocki, & Hawksworth, 1999). RICs are thought to be responsible for the production of the collagenous and noncollagenous extracellular matrices after kidney injury (Whiting et al., 1999). Therefore, RICs play a central role in renal fibrosis. Under normal physiological conditions, RICs are a major site of cyclooxygenase (COX) expression, and thus a major site of COX-derived prostanoid synthesis. Many studies have been performed on the role of RICs in renal function and diseases (Whiting et al., 1999; Zhang et al., 2018). However, the role of RICs during nephron regeneration has not been elucidated.

Prostaglandin E2 (PGE2) is synthesized from arachidonic acid by COX into an intermediate that is metabolized into active PGE2 (Park, Pillinger, & Abramson, 2006). PGE2 has been demonstrated to play an important role in the development of the pronephros (Chambers, Addiego, Flores-Mireles, & Wingert, 2020; Marra et al., 2019; Poureetezadi, Cheng, Chambers, Drummond, & Wingert, 2016). Modulating levels of PGE2 restrict the formation of the distal segment and expanded a proximal segment lineage (Poureetezadi et al., 2016). PGE2 also regulates the development of renal multicellular cells in zebrafish. PGE2-deficient embryos form fewer renal multicellular progenitors and instead develop more transporter cells (Marra et al., 2019). However, whether PGE2 affects nephrogenesis during the development and regeneration of the zebrafish mesonephron has not been investigated.

Here, single-cell sequencing of zebrafish kidney revealed that *fabp10a* was a marker of RICs, and *Tg(fabp10a:GFP)* transgenic line can specifically label RICs in zebrafish kidney. The formation of RICs and nephrons are closely accompanied. COX2 expressing RICs were in close contact with the renal progenitor cell aggregates during nephron regeneration. Inhibiting PGE2 production could inhibited nephrogenesis. PGE2 activated the Wnt signaling pathway in renal progenitor cell aggregates and rescued the Wnt4a deficiency during nephron regeneration. These results suggest that RICs secret PGE2 to promote the maturation of renal progenitor cell aggregates by activating the Wnt signaling pathway.

## Results

### Single-cell messenger RNA sequencing reveals renal interstitial cells

Understanding the function of an organ requires the characterization of all its cell types (Panina, Karagiannis, Kurtz, Stacey, & Fujibuchi, 2020). To illustrate the functions of the various cell types in the zebrafish kidney, we sequenced the kidney cells by single-cell messenger RNA sequencing. Six randomly selected zebrafish kidneys were used and obtained about 7,147 cells for RNA sequencing using a modified version of the cell expression by linear amplification and sequencing (CEL-seq) method and incorporating unique molecular identifiers to count the transcripts. Our initial analysis of the kidney samples identified distinct tSNE clusters that were comprised of 12 different cell types (*Figure 1A*). We identified unique gene expression signatures that defined each of these cell clusters. The identities of the 11 clusters were readily assigned by expressing known markers or previous sequencing data (Tang et al., 2017). For example, two kidney clusters were comprised of epithelial cell types that are highly conserved with mammals, including the distal tubule defined by expression of *solute carrier family 12 member 3 (slc12a3*), *ATPase Na+/K+ transporting subunit alpha 1a, tandem duplicate 4* (*atp1a1a.4*), and *ATPase Na+/K+ transporting subunit beta 1a* (*atp1b1a*), as well as the proximal tubule defined by the expression of *solute carrier family 22 member 2* (*slc22a2*), *paired box 2a* (*pax2a*), and *aminoacylase 1*(*acy1*) (Tang et al., 2017). We also identified hematopoietic stem cells, erythrocytes, T cells, neutrophils, macrophages, and vascular endothelial cells (*Figure 1A, B, Figure 1-source data 1*). Interestingly, we found a new cell type (cluster 12). These cells expressed well-known markers of interstitial cells, including *collagen type 1 alpha 1b* (*col1a1b*), *collagen type 1 alpha 2* (*col1a2*), *collagen type 6 alpha 4a* (*col6a4a*), and *matrix metallopeptidase 2* (*mmp2*) (Fitzgerald, Rich, Zhou, & Hansen, 2008; Guo et al., 2020; Kallakury et al., 2001; Wang et al., 2021) (*Figure 1B and Figure 1*—*figure supplement 1A-D*). We speculate that this (Her, Chiang, Chen, & Wu, 2003) cell type was RICs. To further determine the function of this cell cluster, we analyzed all genes of this cell cluster via a Gene Ontology (GO) analysis. A significant percentage of the genes corresponded to the extracellular matrix, the extracellular region, and the response to steroid hormones were found. This cluster of cells was also enriched in genes that respond to lipids, which are richly contained by RICs in mammals (*Figure 1C*) (Hao & Breyer, 2007). Further analysis the marker of this cell cluster, we also found *the fatty acid-binding protein 10a* (*fabp10a*), which encodes a small protein that plays an important role in intracellular binding and trafficking of long-chain fatty acids (Her et al., 2003), was specifically expressed (*Figure 1D*). So we inferred that *fabp10a* was a specific marker of this cell cluster.

**Figure 1.**
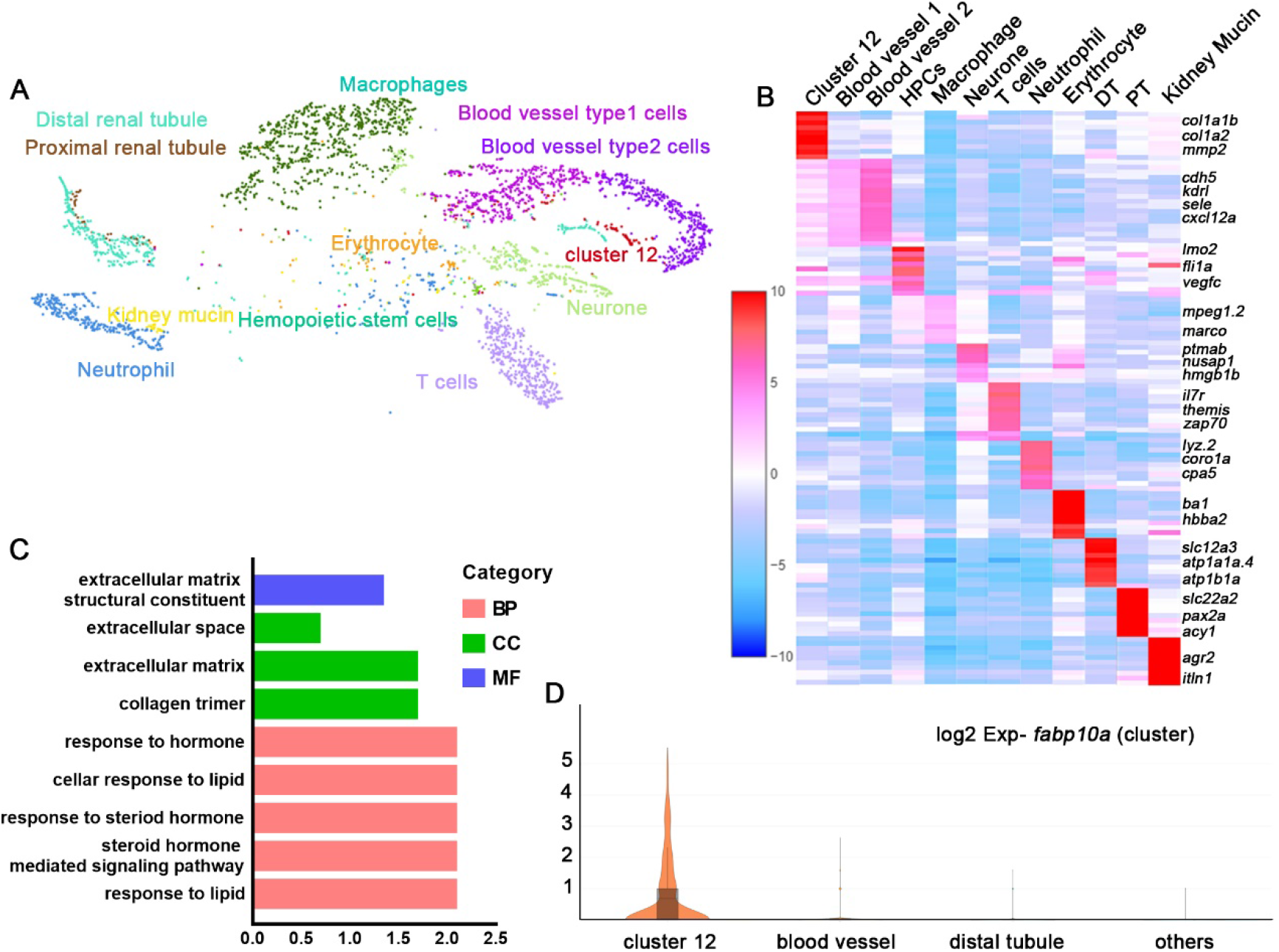
Single messenger RNA sequencing of kidney cells. (**A**) A tSNE plot showing the clustering of all cells of kidney after alignment using the Seurat package. Cells clustered with resolution 0.6. (**B**) Heat map showing relative log-expression of the top 2 or 3 marker genes for each cell cluster identified in Figure 1A. (**C**) GO analysis of differentially expressed genes for cluster 12. False discovery rate <0.05 was considered as significantly enriched. (**D**) Expression analysis of *fabp10a* in all cluster and found that *fabp10a* was specifically expressed in cluster 12. DT, Distal tubule; PT, Proximal tubule; HPCs, Hematopoietic stem cells.

### The *Tg(fabp10a:GFP)* line specifically labels RICs in zebrafish

*Tg(fabp10a:GFP)* has been reported to specifically label the liver during early embryonic development (Her et al., 2003). To verify our conjecture, we crossed *Tg(fabp10a:GFP)* with *Tg(cdh17:DsRed)*, which is a transgenic line specifically labeled with renal tubules, and found that *Tg(fabp10a:GFP)* not only labeled the liver but also a small part of the pronephros at 4 day post-fertilization (4 dpf). After 11 dpf, we found a large number of *Tg(fabp10a:GFP)* labeled cells were distributed around the area where the mesonephric tubules appeared (*Figure 2*—*figure supplement 1A*). These cells were fusiform or irregular polygons, with radiating cytoplasmic projections, and interweaved between cells to form a network. It was difficult to detect the green fluorescent signal in the newly formed mesonephric tubules during this phase (*Figure 2A*). *Tg(fabp10a:GFP)* labeled cells in zebrafish and RICs in mammals have the same shapes and positions (Zhang et al., 2018), so we inferred that the *Tg(fabp:10a:GFP)* transgenic line can be used to specially label RICs in juvenile zebrafish kidney.

**Figure 2.**
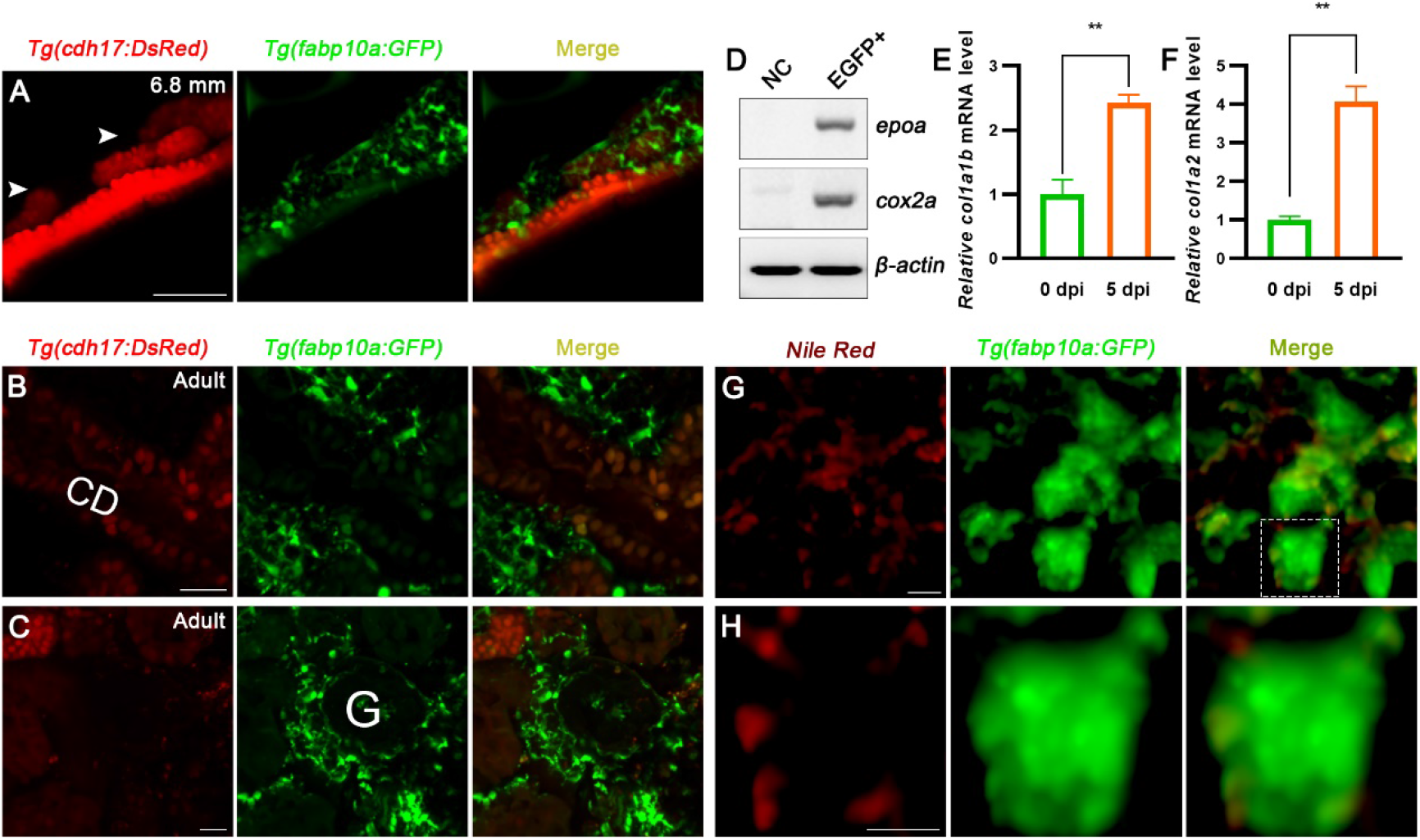
RICs can be specifically labeled by *Tg(fabp10a:GFP)* line. (**A**) The localization of *Tg(fabp10a:GFP)* labeled cells in 6.8 mm stage juvenile zebrafish. *Tg(cdh17:DsRed)* labeled kidney tubules (Arrowheads, new mesonephric branches; n=6). (**B, C**) The localization of *Tg(fabp10a:GFP)* labled cells in adults zebrafish kidney (n=3). These cells tightly wrap kidney tubules (**B**) and glomerulus (**C**) and form network. CD: collecting duct; G: glomerulus. (**D**) PCR analysis the expression of *epoa* and *cox2a*. *β-actin* was used as samples control. GFP^+^ means which cells only have GFP fluorescence and control means all cells except GFP^+^/DsRed^−^ cells. (**E, F**) qPCR relative quantification of *col1a1b* and *col1a2* mRNA in *Tg(fabp10a:GFP)* labeled GFP^+^/DsRed^−^ cells. Both of them was significantly increase at 5 dpi (n=3). Both gene was normalized to the mean expression level at 0 dpi which was set to 1. **P < 0.005 by one-way ANOVA and Dunnett’s post-test. (**G**) Nile red strained the section of *Tg(fabp10a:GFP)* zebrafish kidney showing *Tg(fabp10a:GFP)* labeled cells contained plentiful lipid droplets. **H**: Higher-magnification image of boxed area shown in **G**. Scale bar in A, B, C, 100 μm; G, H, 10 μm.

To further identify these cells, we observed sections of the *Tg(fapb10a:GFP; cdh17:DsRed)* adult kidney. A large number of *Tg(fabp10a:GFP)* transgenic line labeled cells were attached to the renal tubules and collecting ducts, or distributed in the renal interstitium. Moreover, these cells also existed on the lateral side of the glomerulus and tightly surrounded the glomerulus. A few renal tubular epithelial cells were labeled by the *Tg(fabp10a:GFP)* line, but the green fluorescence was weak and hardly detectable. As in juvenile zebrafish, these cells had multitudinous shapes and formed a network (*Figure 2B, C*). The distribution and morphology of these cells were the same as those of the RICs in the mammalian metanephros (Whiting et al., 1999). Next, we sorted the adult kidney cells of the *Tg(fapb10a:GFP; cdh17:DsRed)* double transgenic line by flow cytometry. Two groups of cells were collected. One group of cells only expressed green fluorescence (GFP^+^/DsRed^−^), and the control group included all of the other cells, except GFP^+^/DsRed^−^ cells. We detected the expression of *fabp10a* in GFP^+^/DsRed^−^ cells by semi-quantitative polymerase chain reaction (PCR) analysis, and *fabp10a* was highly expressed in these cells (*Figure 2*—*figure supplement 1B*). This demonstrated that GFP^+^/DsRed^−^ was *fabp10a* positive. It has been reported that EPO is highly expressed as a marker in mammalian RICs (Souma, Suzuki, & Yamamoto, 2015). Using semi-quantitative PCR, we detected *epoa*, the *EPO* homologous gene in zebrafish, and found that *epoa* was expressed only in GFP^+^/DsRed^−^ cells, and almost no expression was detected in the control. COX2 is also considered a marker of RICs (Zhang et al., 2018). *cox2a*, the *COX2* homologous gene in zebrafish, was only expressed in GFP^+^/DsRed^−^ cells (*Figure 2D*). It has been reported that RICs differentiate into fibroblasts when AKI occurs in mammals (Baues et al., 2020). We injected gentamicin to induce AKI in zebrafish and collected the kidney GFP^+^/DsRed^−^ cells of *Tg(fapb10a:GFP; cdh17:DsRed)* at 5 days post-injuring (5 dpi) by flow cytometry. Compared with the uninjured kidney, the fibroblast markers *col1a1b* and *col1a2* mRNA increased in GFP^+^/DsRed^−^ cells 5 dpi (*Figure 2E, F*). Moreover, mammalian RICs are lipid-rich cells (Hao & Breyer, 2007). Therefore, Nile red, a fluorescent dye that specifically stains lipid, was used to detect the lipid droplets in renal tissue sections of adults. As a result, many lipid droplets were detected in the cytoplasm of *Tg(fabp10a:GFP)* labeled cells (*Figure 2G, H*). Therefore, these studies confirmed that this cluster of cells and mammalian RICs have a high degree of conservation based on cell location, morphology, and molecular and biochemical characteristics. Thus, we determined that these cells are the RICs of zebrafish.

### Kidney injury promotes RICs to synthesize and secrete PGE2

Kidney injury in adult zebrafish induced the synchronous formation of many new nephrons marked by *lhx1a* (Diep et al., 2011). The RICs were wrapped around and closely interacted with the *lhx1a^+^* cell aggregates at 4 dpi in the *Tg(fapb10a:GFP; lhx1a:DsRed)* injured kidney. At 5 dpi, RICs and progenitor cell aggregates proliferated simultaneously, suggesting that RICs may participate in the regeneration of nephrons (*Figure 3*—*figure supplement 1A*, *B*). The nephron regeneration transcriptome data showed that *cox2a*, RICs marker gene, increased significantly after kidney injury. We confirmed this result by RT-PCR and found that *cox2a* mRNA increased at 1 dpi, reached the highest level at 5 dpi, and returned to normal at 9 dpi (*Figure 3A*). We also determined the expression of Cox2a by Cox2 antibody immunofluorescence, and Cox2a was expressed in RICs during regeneration (*Figure 3*—*figure supplement 1C*). Interestingly, most of the Cox2a-positive RICs were in close contact with progenitor cell aggregates (*Figure 3B*).

**Figure 3.**
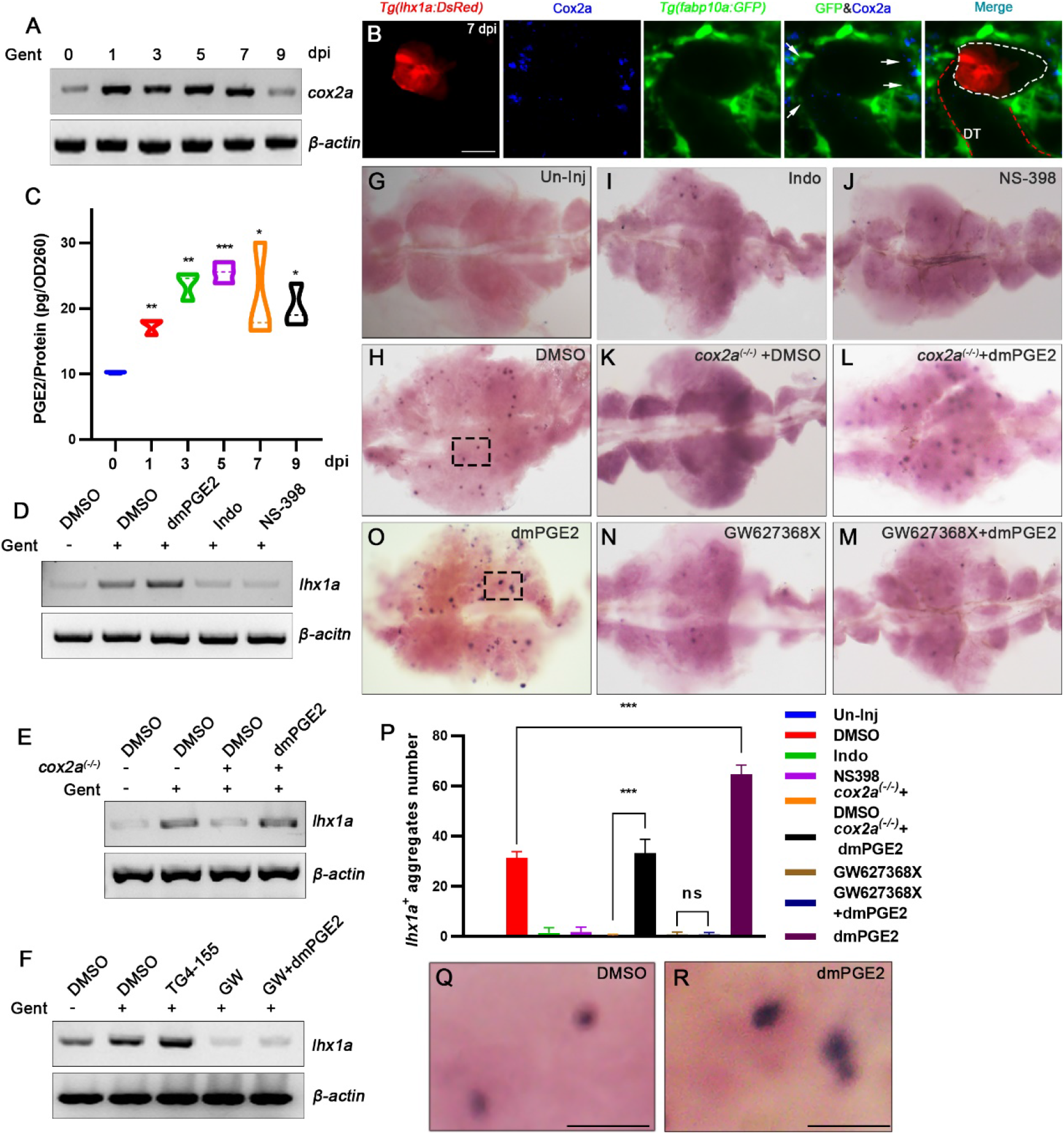
RICs promote the nephron regeneration by secreting PGE2. (**A**) The *cox2a* mRNA level were determined by RT-PCR at 0, 1, 3, 5, 7, 9 dpi. *β-actin* was used as samples control. The result showed that the expression of *cox2a* were up-regulated after AKI and reached the peak levels at 5 dpi. (**B**) Immunofluorescent staining of Cox2a in *Tg(fabp10a:GFP;lhx1a:DsRed)* zebrafish kidney at 7 dpi. RICs around *lhx1a^+^* cell aggregates highly expressed Cox2a (arrows; DT: Distal tubule, red outline; *lhx1a^+^* cell aggregates: white outline, n=4). (**C**) PGE2 level were determined at 0, 1, 3, 5, 7, 9 dpi by PGE2 ELSIA kits. Three individuals were used for each time point. PGE2 levels increased after AKI and reached the peak levels at 5 dpi. Data were analyzed by ANOVA, *p < 0.05, **p < 0.01, ***p < 0.001 versus 0 dpi. (**D**) The *lhx1a* mRNA level were determined by RT-PCR at 7 dpi. The results showed that *lhx1a* mRNA level was up-regulated at 7 dpi, up-regulated when injected dmPGE2 and down-regulated when injected Indo or NS-398. (**E**) The *lhx1a* mRNA level were determined by RT-PCR at 7 dpi in *cox2a^(−/−)^* and wild type zebrafish kidney. (**F**) The *lhx1a* mRNA level were determined by RT-PCR at 7 dpi when injected EP2 inhibitor TG4-155 or EP4 inhibitor GW627368X (GW). *β-actin* was used as samples control in D, E and F. (**G-O**) Whole-mount *lhx1a in situ* hybridization showing the trunk kidney region at 7 dpi (n=5-7). (**G**) Un-injured and DMSO treated kidneys do not have *lhx1a^+^* cell aggregates. (**H**) injury induces the formation of *lhx1a^+^* cell aggregates. Indo (**I**), NS-398 (**J**), or Cox2a deficiency (**K**) inhibit the formation of *lhx1a^+^* cell aggregates. dmPGE2 can rescue the deficiency of Cox2a (**L**). GW627368X inhibits the formation of *lhx1a^+^* cell aggregates (**N**) and dmPGE2 cannot rescue the inhibition (**M**). dmPGE2 can promote the formation of *lhx1a^+^* cell aggregates (**O**). (**P**) *lhx1a^+^* cell aggregates of whole kidney calculated using ImageJ. n= 5-7 fishes for each condition. Data were analyzed by ANOVA, *p < 0.05, **p < 0.01, ***p < 0.001. **Q, R**: The volume of *lhx1a^+^* cell aggregates of injected dmPGE2 kidney (**R**) compared with that of control group (**Q**) (n=5-7). Higher-magnification image of boxed area shown in **O and H**individually. Scale bar in C, 50 μm; Q and R, 500 μm.

Cox2a is a rate-limiting enzyme in PGE2 synthesis (Norregaard, Kwon, & Frokiaer, 2015). Thus, we detected the PGE2 level in the kidney using a PGE2 ELISA kit and found that PGE2 increased significantly during nephron regeneration. PGE2 was about 69.7% higher than that in the uninjured control group at 1 dpi, and reached the highest level at 5 dpi, which was about 2.49 times higher than the control group (*Figure 3C*). This pattern was consistent with the change in *cox2a*. The production of PGE2 is mainly catalyzed by the COX family to produce arachidonic acid and PGH2 and then catalyzed by prostaglandin E synthase (PTGES) to produce PGE2 (Poureetezadi et al., 2016). We determined the expression of the zebrafish homologous genes in the Cox family, *cox1*, and *cox2b*, and found that the expression of *cox1* was not significantly different. *cox2b* was significantly downregulated during nephron regeneration, indicating that only *cox2a* was significantly upregulated during nephron regeneration. We also determined that *ptges* expression was significantly upregulated (*Figure 3*—*figure supplement 1D*-*F*). These results indicate that the increase of PGE2 in injured kidney was caused by RICs specifically expressing *cox2a*, and PGE2 secreted during nephron regeneration was produced by RICs.

### RICs secrete PGE2, which promotes nephron regeneration

These results show that RICs, which surrounded progenitor cell aggregates, highly expressed Cox2a and secreted PGE2, suggesting that PGE2 may play an important role in nephron regeneration. To confirm whether endogenous PGE2 participates in nephron regeneration after AKI, we intraperitoneally injected indomethacin (Indo, 200 μM 10 μL per fish), an inhibitor of both COX1 and COX2 at 2, 4, and 6 dpi (Poureetezadi et al., 2016). As results, *lhx1a* expression was downregulated and *lhx1a^+^* cell aggregates decreased at 7 dpi (1.5 ± 2.1 per kidney; n=7) compared with the control (31.2 ± 2.7 per kidney; n=6) according to *in situ* hybridization (*Figure 3D, G-I, P*). These results suggest that suppressing COX activity restrained nephron regeneration. *cox1* expression did not change significantly during regeneration, whereas the expression of *cox2a* was significantly upregulated. We inferred that *cox2a* plays a decisive role in PGE2 synthesis. N-(2-cyclohexyloxy-4-nitrophenyl) methane sulfonamide (NS-398, 140 μM, 10 μL per fish), a selective COX2 inhibitor (Marra et al., 2019), was intraperitoneally injected after AKI. As a result, *lhx1a* expression was downregulated, as in the Indo group. The numbers of *lhx1a^+^* cell aggregates decreased after the Indo treatment (1.8 ±1.9 per kidney; n=7) (*Figure 3D, G, H, J, P*).

We next utilized a genetic model of Cox2a deficiency to confirm the effect of the Cox2a-PGE2 pathway on nephron regeneration. No obvious differences were detected between *cox2a^(−/−)^* and normal fish, including the size and structure of the adult kidney. To determine the effect of Cox2a deficiency, we injured the kidney of *cox2a^(−/−)^*, and the *lhx1a* mRNA and *lhx1a^+^* cell aggregates (0.3 ± 0.5 per kidney; n=7) decreased compared with the wild type, suggesting that Cox2a is needed for nephrons regeneration. To further confirm that the Cox2a deficiency affecting nephron regeneration was related to PGE2 synthesis, we intraperitoneally injected the stable PGE2 analog dmPGE2 (600 μM, 10 μL per fish) in *cox2a^(−/−)^* (Goessling et al., 2009; Marra et al., 2019), and found that the expression of *lhx1a* and the number of *lhx1a*^+^ cell aggregates (33.1 ± 5.6 per kidney; n=7) was rescued to the level of the wild-type at 7 dpi (*Figure 3E, H, K, L,P*). These results indicate that Cox2a is needed to nephrons regeneration for PGE2 synthesis. To detect the effect of excess PGE2 on renal regeneration, we intraperitoneally injected dmPGE2 (600 μM, 10 μL per fish) after AKI and found that the expression of *lhx1a* and *lhx1a^+^* cell aggregates (64.7 ± 3.6 per kidney; n=7) increased significantly compared with the control group at 7 dpi, suggesting that excessive PGE2 accelerates nephron regeneration (*Figure 3D, H, O, P*). Taken together, these results show that PGE2 synthesized by RICs is necessary to nephrons regeneration.

### PGE2 facilitates nephron regeneration through the EP4 receptor

PGE2 signals through four prostaglandin E receptors (Ptger1–4; EP1-4) in mammals (Miyoshi et al., 2017). In zebrafish, each prostaglandin E receptor has one or more isoforms (Baker & Van Der Kraak, 2019). We sorted the kidney cells of the *Tg(lhx1a:DsRed)* transgenic line by flow cytometry at 5 dpi and obtained DsRed^+^ progenitor cells. Only *ep2a* and *ep4b* mRNAs, but not *ep1*, *ep3*, or *ep4a* mRNAs, were highly expressed in these cells, and the level of *ep4b* mRNA was about twice that of *ep2a* (*Figure 2—figure supplement 2A*). We also determined *ep4b* expression by *in situ* hybridization and confirmed that *ep4b* was expressed in the progenitor cell aggregates (*Figure 3—figure supplement 2B*). To determine which prostaglandin E receptor mediated the effects of PGE2 on the *lhx1a^+^* cell aggregates, a specific pharmacological inhibitor for EP2 (TG4-155, 400 μM, 10 μL per fish) or EP4 (GW627368X, 200 μM, 10 μL per fish) was intraperitoneally injected at 2, 4, and 6 dpi (Baker & Van Der Kraak, 2019). As results, only the EP4 inhibitor GW627368X blocked nephron regeneration (0.8 ± 1.0 per kidney; n=7). In addition, dmPGE2 did not rescue the inhibitory effect on nephron regeneration induced by these inhibitors (*Figure 3F, H, M, N, P*; 0.8 ± 0.8 per kidney; n=7). This result verifies that PGE2 promotes nephron regeneration through EP4, but not other receptors.

### PGE2-EP4 signaling promotes the proliferation of nephron progenitor cell aggregates

The volume of *lhx1a^+^* progenitor cell aggregates increased significantly in the dmPGE2 treated group compared to the control (*Figure 3Q, R*). So, we speculated that PGE2 may play a role in affecting the expansion of progenitor cell aggregates. To confirm this hypothesis, we assessed the proliferation of nephron progenitor cells by EdU assay. First, we blocked PGE2 synthesis with Indo or NS-398 after AKI and assayed the EdU incorporation associated with DsRed^+^ cell aggregates in *Tg(lhx1a:DsRed)* zebrafish kidney. The results showed that most of the progenitor cell aggregates remained diminutive and quiescent in the treatment group, and EdU was rarely detected in these cells (only about 4.2% (Indo; n=7), 3.9% (NS-398; n=7) of aggregates showing 5 or more EdU^+^ nuclei; *Figure 3*—*figure supplement 1G-J*). However, a large number of EdU^+^ DsRed^+^ cell aggregates were found in the controls (intraperitoneally injected DSMO) at 5 dpi, which occurred in an elongating immature nephron tubular structure (about 33.2% of *lhx1a^+^* cell aggregates contained more than five EdU^+^ nuclei; n=6) (*Figure 4A, B*). We also carried out the EdU assay in *cox2a^(−/−)^* and the injured *Tg(lhx1a:DsRed)/cox2a^(−/−)^* zebrafish kidney at 5 dpi. The *lhx1a^+^* cell aggregates in *cox2a^(−/−)^* were also diminutive and quiescent, and only 1.4% were labeled with EdU (n=7). Then, we injected dmPGE2 to rescue the Cox2a deficiency after AKI, and a large number of EdU^+^ DsRed^+^ cell aggregates was detected at 5 dpi (about 35.2% of *lhx1a^+^* cell aggregates contained more than 5 EdU^+^ nuclei; n=7) (*Figure 4C-F*). These results indicate that the PGE2 secreting RICs are needed for the proliferation of progenitor cell aggregates.

**Figure 4.**
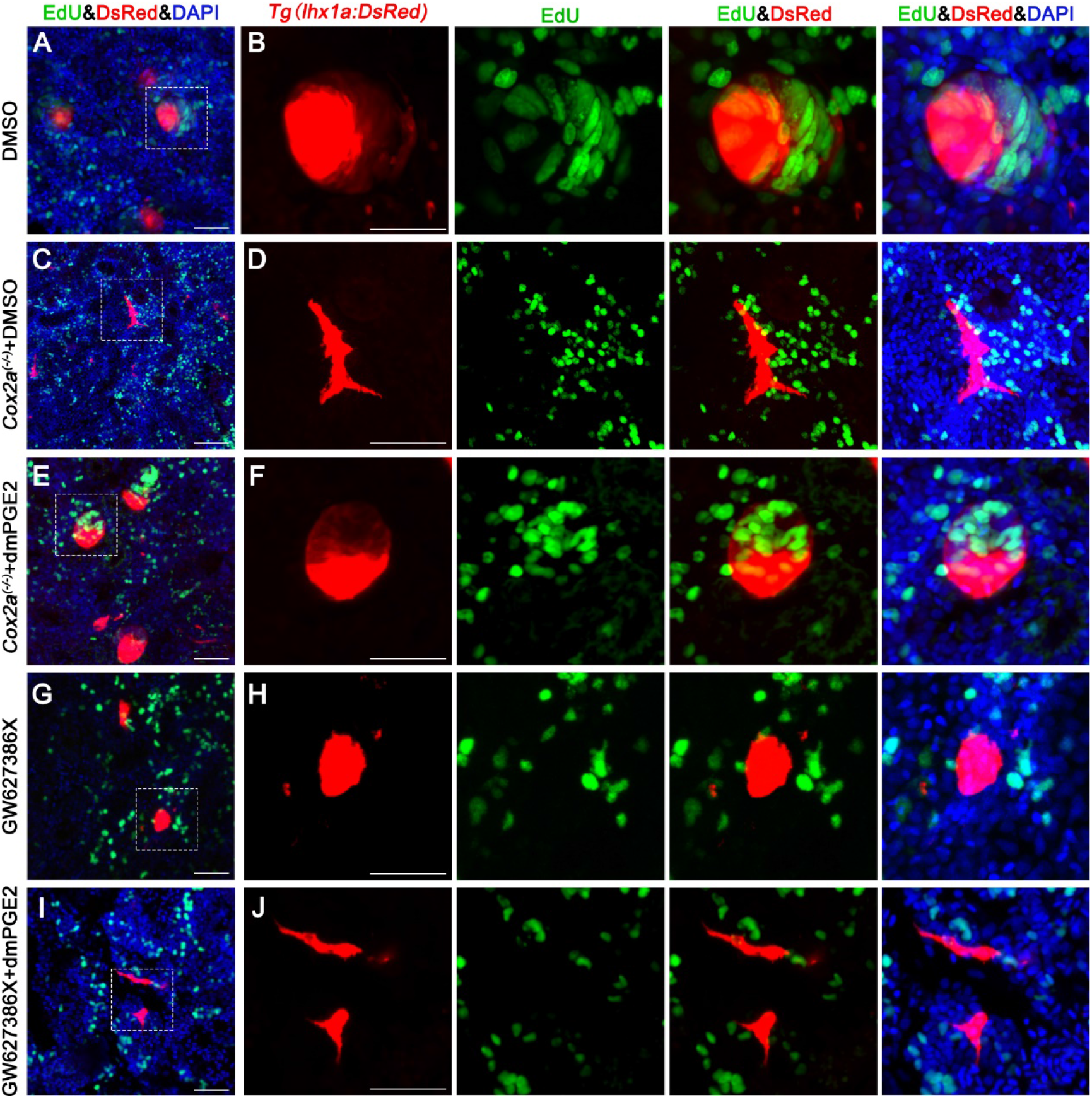
PGE2-EP4 signaling promote the proliferation of progenitor cell aggregates. *Tg(lhx1a:DsRed)* zebrafish were injured by gentamicin injection, injected with EdU to label proliferating nuclei at 5 dpi and kidneys were harvested 3 hours later. (**A, B**) Gentamicin induces *lhx1a^+^* newborn nephrons with proliferating EdU^+^ nuclei. (**C, D**) *cox2a* deficiency reduces the proliferation of *lhx1a^+^* cell aggregates. (**E, F**) dmPGE2 can rescue the proliferation of *lhx1a^+^* cell aggregates in *cox2a* mutant. GW627386X (**G, H**) reduce the proliferation of *lhx1a^+^* cell aggregates and dmPGE2 could not rescue the proliferation of GW627386X treatment group (**I, J**). N= 4-6 fish for each condition. Scale bar, 50 μm. Right images showed the higher-magnification image of boxed area shown in left.

We also blocked the EP4 receptor using GW627368X, and detected the same phenomenon as for *cox2a^(−/−)^* (only about 2.3 % of aggregates showing five or more EdU^+^ nuclei). We tried to rescue this blockade by intraperitoneally injecting dmPGE2, but dmPGE2 did not rescue blocking of the EP4 receptor (only 2.5% of aggregates had five or more EdU^+^ nuclei) (*Figure 4G-J*). These results suggest that PGE2 promotes the proliferation of nephron progenitor cells through the EP4 receptor. Cell proliferation in other kidney cells was not inhibited by Indo, NS-398, Cox2a deficiency, or GW627368X, suggesting that inhibiting the synthesis of PGE2 or PGE2-EP4 signaling specifically suppresses the proliferation of progenitor cell aggregates.

### PGE2-EP4 promotes nephron regeneration through Wnt signaling

Previous reports have shown that PGE2 plays a role in the development of hematopoietic stem cells and liver regeneration by regulating the Wnt signaling pathway(Goessling et al., 2009; North et al., 2010). To determine whether PGE2-EP4 also plays a role in renal regeneration through Wnt signaling, we tested Wnt activity at 7 dpi through the expression of *lef1*, a canonical marker of Wnt signal activity (Kamei, Gallegos, Liu, Hukriede, & Drummond, 2019), and found that Wnt activity increased significantly in the injured kidney compared to the controls. However, Wnt activity was significantly diminished in *cox2a^(−/−)^* than the wild type at 7 dpi, and was completely rescued by intraperitoneally injecting dmPGE2 (*Figure 5A*). Wnt activity was also significantly inhibited by intraperitoneally injecting Indo, NS-398, and the EP4 specific inhibitor GW627368X, but was significantly enhanced by intraperitoneally injecting excessive dmPGE2 (*Figure 5—figure supplement 1A*). We also tested the Wnt activity of the progenitor cell aggregates by β-catenin immunofluorescence at 5 dpi, and showed that most of the β-catenin localized in *lhx1a^+^* cell aggregates and the remainder of the β-catenin was localized in the nucleus (Goessling et al., 2009). β-catenin in the *lhx1a^+^* cell aggregates decreased significantly in *cox2a^(−/−)^* and in the Indo, NS-398, and GW627368X treated kidneys compared to the control at 5 dpi. The effect of Cox2a deficiency was fully rescued by intraperitoneally injecting dmPGE2 (Figure 5*B*-*D*, *Figure 5—figure supplement 1B*-*E*). These results show that PGE2-EP4 signaling regulates Wnt activity in *lhx1a^+^* cell aggregates by regulating the stability and localization of β-catenin during nephron regeneration. To further verify this relationship, we intraperitoneally injected the Wnt signaling inhibitor XAV939 (200 μM, 10 μL per fish), to promote degradation of β-catenin after AKI (Hofsteen et al., 2018). As a result, the regeneration of nephrons was inhibited at 7 dpi and was rescued by dmPGE2 (*Figure 5G-J, M, Figure 5—figure supplement F*).

**Figure 5.**
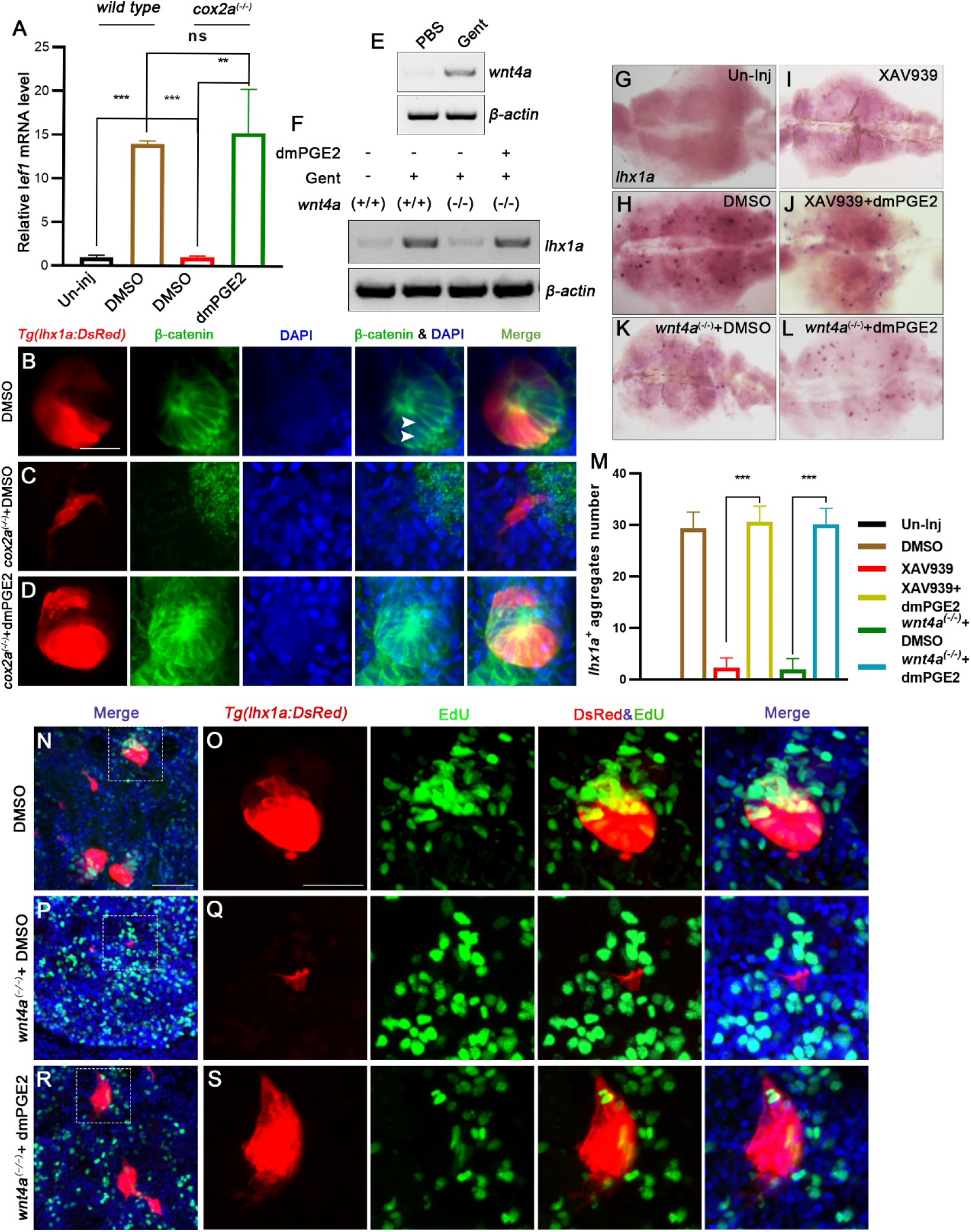
PGE2-EP4 signaling promote nephron regeneration through Wnt signaling. (**A**) qPCR relative quantification of *lef1* mRNA in kidney tissue of wild type and *cox2a* mutant harvested 7 dpi. Cox2a deficiency can decrease *lef1* level and intraperitoneal injection of dmPGE2 can rescue the influence of Cox2a deficiency. Gene was normalized to the mean expression level of un-injured kidney which was set to 1. Data were analyzed by ANOVA, **p < 0.005, ***p < 0.001, ns, no significant difference (n = 3 independent experiments). (**B-D**) Immunofluorescent staining of β-catenin in *Tg(lhx1a:DsRed)* zebrafish kidney at 5 dpi. (**B**) Injected DSMO as control group and amount of β-catenin can be detected in *lhx1a^+^* cell aggregates cytoplasm and nucleus (arrowheads). Cox2a deficiency (**C**) decreases β-catenin level in *lhx1a^+^* cell aggregates. and intraperitoneal injection of dmPGE2 (**D**) can rescue the influence of Cox2a deficiency. (**E**) The *wnt4a* mRNA level were determined by RT-PCR in un-injured or injured kidney. *β-actin* was used as samples control. *wnt4a* mRNA increased at 7 dpi. (**F**) The *lhx1a* mRNA level were determined by RT-PCR at 7 dpi in *wnt4a^(−/−)^* and wild type zebrafish kidney. *β-actin* was used as samples control. (**G-L**) Whole-mount *lhx1a in situ* hybridization showing the trunk kidney region at 7 dpi. Injection of XAV939 (**I**) or Wnt4a deficiency (**K**) can reduce the *lhx1a^+^* cell aggregates and dmPGE2 can rescue the influence of XAV939 treatment (**J**) or Wnt4a deficiency (**L**) (n=5-7 in each condition). (**M**) *lhx1a^+^* cell aggregates of whole kidney calculated using ImageJ. N=5-7 fishes for each condition. Data were analyzed by ANOVA, ***p < 0.001. (**N, O**) Gentamicin induces *lhx1a^+^* new nephrons with proliferating EdU^+^ nuclei (n=4). (**P, Q**) Wnt4a deficiency reduces the proliferation of *lhx1a^+^* cell aggregates (n=4). (**R, S**) dmPGE2 can rescue the Wnt4a deficiency and recovered the proliferation of *lhx1a^+^* cell aggregates in *wnt4a* mutant (n=4). O, Q and S showed the higher-magnification image of boxed area shown in N, P and R. Scale bar in B-D, 50 μm; N-S 100 μm.

Genetic studies in mice have shown that canonical Wnt signaling plays various roles in kidney development and *Wnt4* expression in the nephrogenic cap mesenchyme and plays central roles in the renal progenitor cell aggregate expansion stages of nephrogenesis (Kispert, Vainio, & McMahon, 1998; Pulkkinen, Murugan, & Vainio, 2008). We sorted *lhx1a^+^* cells by flow cytometry using the *Tg(lhx1a:DsRed)* zebrafish kidney at 5 dpi, and found that the *Wnt4* homologous gene in zebrafish *wnt4a* was expressed in these cells. We also determined that the level of *wnt4a* mRNA increased at 7 dpi (*Figure 5E, Figure 5—figure supplement 1G*). Therefore, Wnt4a may play a role in the regeneration of nephrons. We injured the kidney of *wnt4a^(−/−)^* and discovered that the expression of *lhx1a* and *lhx1a^+^* cell aggregates decreased significantly compared to the wild type at 7 dpi. The deficiency of Wnt4a was fully rescued by an intraperitoneal injection of dmPGE2 (*Figure 5F-M*).

Next, we detected the proliferation of progenitor cell aggregates using the EdU assay in *wnt4a^(−/−)^* and XAV939 treated kidneys. As a result, the proliferation of progenitor cell aggregates decreased significantly at 5 dpi. Only 4.2% (intraperitoneally injected XAV939) and 3.5% (Wnt4a deficiency) of aggregates showing 5 or more EdU^+^. An intraperitoneal injection of dmPGE2 eliminated the effect of Wnt4 deficiency and the XAV939 treatment, and a large number of EdU^+^ DsRed^+^ cell aggregates were detected. Approximately 30.2 % of the *lhx1a^+^* cell aggregates contained more than 5 EdU^+^ nuclei in both group and no significant difference was detected from the control (which was intraperitoneally injected with DMSO) (*Figure 5N-S, Figure 5-figure supplement 1H-M*). These results suggest that PGE2-EP4 signaling affects the expansion of progenitor cell aggregates by regulating the Wnt signaling pathway. As PGE2 and Wnt4a affect the stability of β-catenin through independent pathways. And PGE2 or Wnt4a alone could not effectively activate the Wnt signaling pathway during regeneration. This indicates that activation of the Wnt signaling pathway during nephron regeneration requires the cooperation of multiple factors.

### Real-time observations of the interaction between RICs and renal progenitor cells

Because our experiments were mainly conducted in adult fish, we could not determine the real-time interaction between RICs and renal progenitor cells. However, zebrafish are transparent during the mesonephrogenesis stage and were used to achieve this objective (Diep et al., 2011; Jerman & Sun, 2017). During development of the mesonephron from 10 dpf to 16 dpf, the *Tg(fabp10a:GFP; cdh17:DsRed)* and *Tg(fabp10a:GFP; lhx1a:DsRed)* fish were observed. Renal progenitor cells occurred with the pronephric tubules at the 4.6 mm stage. The RICs were detected around the progenitor cells at the 5 mm stage, and the progenitor cell aggregate as a progenitor cell aggregate. More RICs appeared and covered the cell aggregates as a cell network. The progenitor cell aggregates elongated to form the first mesonephron branch at the 5.3 mm stage, and large numbers of RICs appeared in the direction that the progenitor cell aggregates elongated. The mesonephric tubule elongated into the nascent nephron at the 5.8 mm stage, and a large number of RICs formed a network that completely wrapped the entire nephron. The distribution of RICs was consistent with adult zebrafish (*Figure 6A-F, Figure 6*—*movie supplement 1*). As the body grew, other nascent nephrons appeared in the pronephric tubules. RICs repeated the above process to produce a new cell network to wrap nascent nephrons (*Figure 2A*). This finding demonstrates that the RIC network is also a basic element of nephron structure. At the same time, we determined the proliferation of RICs during this process. A large number of RICs were EdU^+^ as in regeneration, indicating that cell proliferation during nephrogenesis is a source of RICs (*Figure 6G, Figure 3*—*figure supplement 1B*).

**Figure 6.**
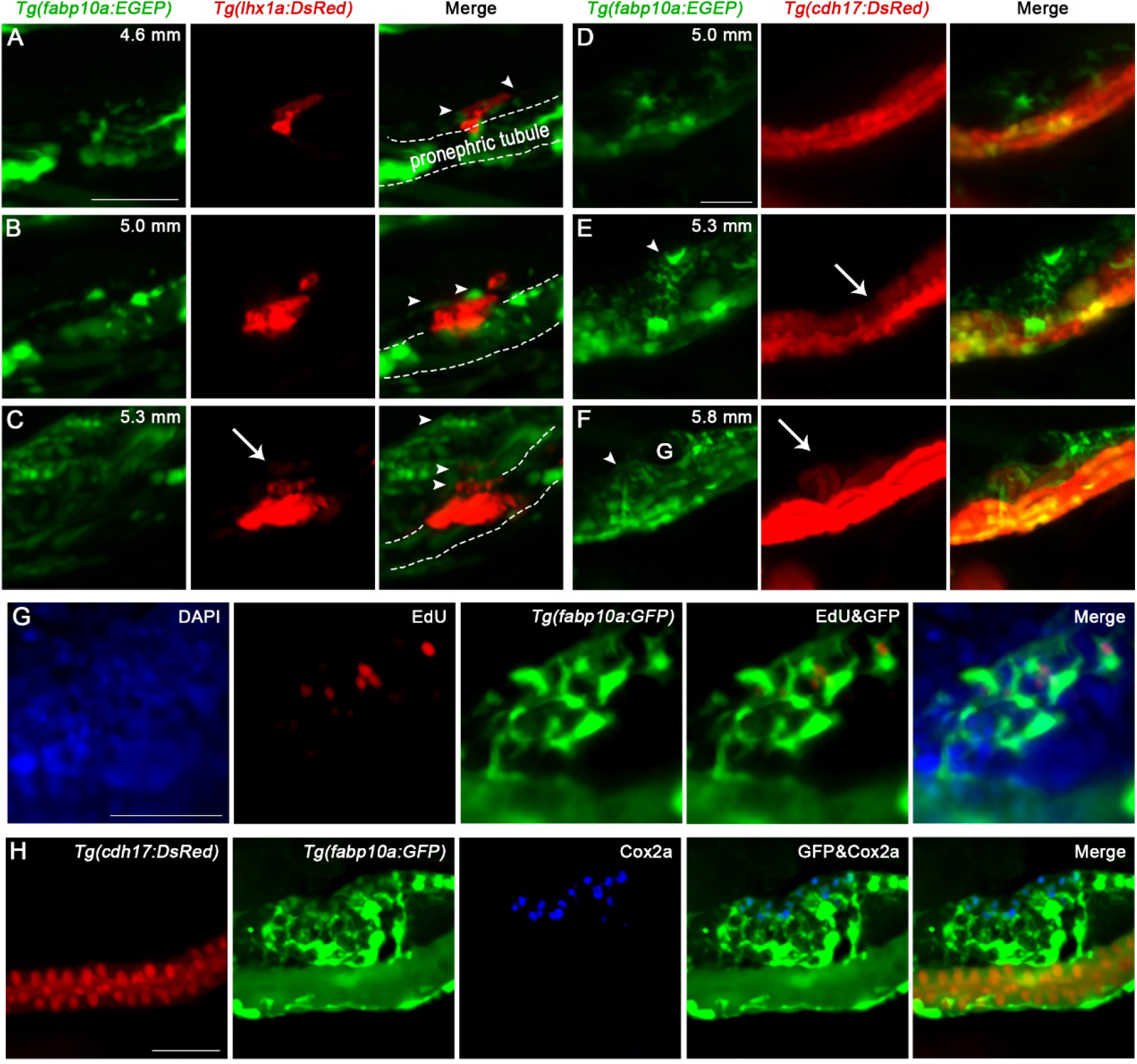
The interaction between RICs and renal progenitor cells. (**A-C**) Time-lapse microscopy of *Tg(fabp10a:GFP;lhx1a:DsRed)* reveals the interaction between RICs (arrowheads) and progenitor cell aggregate at 4.6 mm (**A**), 5.0 mm (**B**) and 5.3 mm (**C**) stage. (**D-E**) Time-lapse microscopy of *Tg(fabp10a:GFP;cdh17:DsRed)* reveals the behavior of RICs during mesonephric development (n=4). (**D**) The RICs (arrowheads) occur upon the pronephric tubule at 5.0 mm stage (n=4). (**E**) Mesonephric branch (arrow) occur upon the pronephric tubule and be wrapped by RICs (arrowheads) at 5.3 mm stage (n=4). (**F**) The mesonephric tubule elongates into nascent nephron (arrow) and RICs (arrowheads) form a network that completely wrap up the tubule at 5.8 mm stage (n=4). (**G**) EdU assay of RICs shown that the RICs were proliferating during mesonephric development (n=5). (**H**) Immunofluorescent staining of Cox2a in *Tg(fabp10a:GFP;cdh17:DsRed)* zebrafish at 5.2 mm stage shown that Cox2a was specifically expressed in RICs (arrow, n=5). Scale bar in A-C, 50 μm; B-D, 100 μm; G, H, 50 μm.

Furthermore, we detected Cox2a expression in the RICs of juvenile fish and found that Cox2a was specifically expressed in RICs adjacent to the progenitor cell aggregates as in the adult kidney (*Figure 6H, Figure 3B*). This result infers that Cox2a is also required for the development of the mesonephron. To confirm this inference, we employed *Tg(cdh17:Dendra2-NTR)/cox2a^(−/−)^* fish. The branch of the first mesonephron was not observed under the pronephric tubules at the 5.3 mm stage but was easily found in the DSMO treated group at this stage. We also treated *Tg(cdh17:Dendra2-NTR)* with dmPGE2 and found that elongation of the mesonephron was accelerated by dmPGE2 (*Figure 6*—*figure supplement 1*). These results suggest that RIC-secreted PGE2 promotes nephrogenesis and the development of the mesonephron.

## Discussion

RICs can be labeled in mice with the FoxD1-Cre and tenascin-c-creer2 transgenic lines (He et al., 2013; Humphreys et al., 2010). However, these lines have not been reported in zebrafish. Therefore, new markers are needed for zebrafish RICs. Our results of single-cell sequencing of the zebrafish kidney showed that *fabp10a* can be used as a RIC marker. Subsequently, we proved that RICs were specifically labeled by the *Tg(fabp10a:GFP)* transgenic line. RICs transgenic zebrafish lines allow for fluorescence-activated cell sorting and analysis of RICs, *in vivo* imaging of the interactions between RICs and other cells, and RIC-focused chemical genetic screens. Therefore, the RIC transgenic line is a major step forward in the study of RIC behavior and functions *in vivo*.

Taking advantage of real-time observations in zebrafish juveniles, we examined the interactions between RICs and renal progenitor cells. We discovered that RICs form a network wrapping the entire cell aggregate. COX2 was highly expressed during the regeneration of nephrons in RICs, which were in close contact with the cell aggregates, and suggested that RICs directly secrete PGE2 that acts on the cell aggregates. Most of the renal progenitor cell aggregates remained in the early state and did not proliferate and differentiate normally when PGE2 production was inhibited. Moreover, the expression of *col1a1b* and *col1a2* in RICs increased significantly during regeneration, proved that RICs secreted a large amount of extracellular matrix, which is also a crucial part of the microenvironment of progenitor and stem cells (Watt & Huck, 2013). These results indicate that RICs provide a necessary microenvironment for the renal progenitor cells through the formation of a cell network. It also demonstrates that PGE2 is a crucial signaling molecule to rapidly induce the formation of new nephrons. The mechanism that promotes the secretion of PGE2 by RICs will be an interesting topic for further research.

Furthermore, our results show that the PGE2 receptor EP4b was highly expressed in renal progenitor cells. A specific inhibitor treatment revealed that PGE2 functioned through the EP4 receptor during nephron regeneration. EP4 modified Wnt signaling at the level of β-catenin degradation through cAMP/PKA-mediated stabilizing phosphorylation events (Goessling et al., 2009). We further confirmed that PGE2 activates the Wnt signaling pathway in renal progenitor cell aggregates. Wnt4 is a key auto-regulator of the mesenchymal to epithelial transformation during nephrogenesis in the mouse kidney (Kispert et al., 1998). The zebrafish *wnt4a* mutant has the same defect as PGE2 deficiency during nephron regeneration, and the nephron cannot differentiate and mature normally. Injecting dmPGE2 rescued the Wnt4a deficient phenotype. Unlike PGE2, Wnt4a activates the Wnt signaling pathway by binding the frizzled receptor and the low-density lipoprotein co-receptor (Yang et al., 2020). Deleting PGE2 or Wnt4a alone will cause a defect in activation of Wnt signaling, demonstrating that activating the Wnt signal requires the cooperation of multiple factors during nephrogenesis. As PGE2 is a small metabolite, it can be used to promote nephron regeneration and in vitro kidney organoids culture soon.

Here, RICs formed a cellular network that wrapped the renal progenitor cell aggregates during nephron development and regeneration. The RICs promoted the proliferation and maturation of nephrons by secreting PGE2. In general, RICs provide a microenvironment for the differentiation of renal progenitor cells, which is necessary for the rapid recovery of damaged nephrons. Our research not only enriches the cell interactive mechanisms of nephrogenesis but also determined that the RICs cell network develops as a basic element of each nephron, which lays a foundation for the development of new methods for nephron regeneration and replacement.

## Materials and Methods

### Zebrafish

Zebrafish embryos, larvae, and adults were produced, grown and maintained according to standard protocols described in the zebrafish book. For experiments with adult zebrafish, animals ranging in the age from 3 to 12 months were used. Zebrafish were maintained in standard conditions under a 14-hours-light and 10-hours-dark cycle and fed three times daily. Zebrafish were anesthetized using 0.0168% buffered tricaine (MS-222, Sigma). AB strains of zebrafish were used as wild-type for this study. The following zebrafish lines were used in this study: *Tg(fabp10a:GFP)* (Her et al., 2003); *cox2a^(−/−)^* (Li, Jin, & Zhong, 2019); *wnt4a^(−/−)^* were purchased from the Chinese National Zebrafish Resource Center (Wuhan, China); *Tg(cdh17:DsRed)*, *Tg(cdh17:dendra2-NTR) and Tg(lhx1a:DsRed)* were constructed in this study.

### Generation of *Tg(lhx1a:DsRed)* transgenic line

We obtained the *lhx1a:EGFP*/pI-SceI plasmid from Dr. Neil A Hukdiede (Swanhart et al., 2010), substituted DsRed for EGFP using ClonExpress II One Step Cloning Kit (Vazyme, C112-01), and constructed *lhx1a:DsRed*/pI-SceI plasmid. 30 pg of *lhx1a:DsRed*/pI-SceI plasmid DNA was injected into 1-cell stage embryos along with the I-SceI restriction enzyme (NEB, R0694S). These injected embryos were raised to adulthood and screened for DsRed expression in known *lhx1a* expression domains and isolated a stable *Tg(lhx1a:DsRed)* transgenic line.

### Generation of *Tg(cdh17:DsRed)* and *Tg(cdh17:Dendra2-NTR)* transgenic line

Promoters of *cadherin-17* (*cdh17*) (−4.3k) were amplified from zebrafish genomic DNA by PCR and constructed *cdh17:DsRed*/pI-SceI or *cdh17:Dendra2-NTR*/pI-SceI plasmids. These injected embryos were raised to adulthood and screened for DsRed or Dendra2 expression in known *chd17* expression domains and isolated stable *Tg(cdh17:DsRed)* or *Tg(cdh17:Dendra2-NTR)* transgenic line.

### Zebrafish AKI model

Intraperitoneal injection of gentamicin induced acute kidney injury as previously described (Chen et al., 2019). In brief, Gentamicin (2.7 μg/μL, 20μL per fish) diluted in water was intraperitoneally injected in wild type or other zebrafish lines. And then each injected zebrafish dropped into an individual container. The fish excreting proteinuria at 1 dpi were used to the following study.

### Single cell RNA sequencing

Kidney cells from 6 randomly selected zebrafish was loaded into Chromium microfluidic chips with 30 v chemistry and barcoded with a 10× Chromium Controller (10X Genomics). RNA from the barcoded cells was subsequently reverse-transcribed and sequencing libraries constructed with reagents from a Chromium Single Cell 30 v3 reagent kit (10X Genomics) according to the manufacturer’s instructions. Sequencing was performed with Illumina (NovaSeq) according to the manufacturer’s instructions (Illumina). The Seurat package was used to normalize data, dimensionality reduction, clustering, differential expression. we used Seurat alignment method canonical correlation analysis (CCA) (Butler, Hoffman, Smibert, Papalexi, & Satija, 2018) for integrated analysis of datasets. For clustering, highly variable genes were selected and the principal components based on those genes used to build a graph, which was segmented with a resolution of 0.6. Gene Ontology (GO) enrichment analysis of marker genes was implemented by the cluster Profiler R package, in which gene length bias was corrected. GO terms with corrected P-value less than 0.05 were considered significantly enriched by marker gene.

### Enzyme-Linked Immunosorbent Assay (ELISA) for PGE2

ELISA for PGE2 was conducted using the Prostaglandin E2 ELISA Kit (D751014-0048, BBI) according to manufacturer’s instructions. Briefly, kidneys were collected from 20 weight-matched fishes, washed in ice-cold PBS, and homogenized in 500 μL PBS buffer. Homogenate was spun down at 12000 rpm for 10 min at 4 °C to eliminate particulate, and supernatant collected for ELISA. Assays were run in technical triplicate.

### Inhibitor treatment

During the stage of nephron regeneration, the Indomethacin (I7378-5G, sigma; 400 μM, 10 μL per fish), NS-398(N194-5MG, sigma; 200 μM, 10 μL per fish), TG4-155(S6793, Selleck; 400 μM, 10 μL per fish), dmPGE2 (D0160, sigma. 600 μM, 10 μL per fish), GW627368X (T1978, TOPSCIENCE; 200 μM, 10 μL per fish), XAV939 (S1180, Selleck; 200 μM, 10 μL per fish) was intraperitoneally injected into zebrafish at 2, 4 and 6 dpi. Kidneys was collected at 7 dpi for RNA extraction, *in situ* hybridization or immunofluorescence. For 5 dpi test, inhibitors were intraperitoneally injected at 2 and 4 dpi and samples were collected at 5 dpi. 1% DMSO was applied as their control groups with same conditions. For larva experiment, dmPGE2 (2 μM in egg water) was used to test the influence to mesonephron development. 0.2% DMSO was applied as their control groups with same conditions.

### *In situ* hybridization

*In situ* hybridization was performed as previously described using *lhx1a* and *ep4b* probes (Chen et al., 2019). Briefly, fish with internal organs and hand removed were fixed overnight in 4% paraformaldehyde (PFA). Fixed kidneys were removed from body and permeabilized with proteinase K (10 μg/mL, Roche) in PBT for 1 h with rocking. Digoxigenin-labeled riboprobes were generated from cDNA fragment comprising the sequences of zebrafish *lhx1a* or *LNA-ep4b probe*. Hybridization was performed as described (Chen et al., 2019). Anti-DIG AP antibody and NBT/BCIP substrate (Roche) were used to detect the probe. After color reaction, the images were taken by BX3-CBH microscope (Olympus, Japan).

### EdU ASSAY

Click-iT Plus EDU Alexa Fluor 647 Imaging Kit (C10640, Invitrogen) were used to detect cell proliferation in sections of kidney. Briefly, EdU solution (200 mM, 10 μL per fish) was intraperitoneally injected into fish. After 3 hours, the kidneys were obtained and performed measurement of proliferation.

### Semi-quantitative RT-PCR and quantitative PCR

RNA was isolated from kidney tissue using TRIzol reagent (Invitrogen,15596018) according to the manufacture’s protocol. Prime Script Ⅱ 1st strand cDNA Synthesis Kit (Takara, 9767) was used to synthesize cDNA, which then subjected to PCR using Taq Master Mix (Vazyme, p112-01) for RT-PCR or TB Green Premix EX Taq Ⅱ (Takara, RR820A) for quanti tative PCR. The following primers were used: primers were *lhx1a* (F: GA CAGGTTTCTCCTTAATGTTC, R: CTTTCAGTGTCTCCAGTTGC); *cox2a*, (F: CGCTATATCCTGTTGTCAAGG, R: GATGGTCTCACCAATCAGG); *col 1a1b*, (F: GGTTCTGCTGGTAACGATGG, R: CCAGGCATTCCAATAAGAC C); *col1a2*, (F: CTGGTAAAGATGGTTCAAATGG, R: CACCTCGTAATCCTTGGCT); *lef1* (Kamei et al., 2019); *cox1*, *cox2b and ptges* (FitzSimons e t al., 2020). *The* primers for *ep1a*, (QF: AAATGTCACCTCGAGCAGAC, QR: ACAGGAGAAAGGCCTTGGAT); *ep4b* (QF: ATCGTTCTCATAGCCACGTCCACT, QR: CCGGGTTTGGTCTTGCTGATGAAT), other eps primers were as described (Chen et al., 2019). The *β-actin* (Chen et al., 2019) or *rp13a* (FitzSimons et al., 2020) were used to standard samples.

### Flow cytometry

To obtain the RNA libraries of *lhx1a^+^* cells or RICs, Kidney cells of *Tg(lhx1a:DsRed)* or *Tg(fabp10a:GFP;cdh17:DsRed)* from 10 fishes was manually dissected in 1% PBS and 0.005% Trypsin-EDTA solution at 0 or 5 dpi. Interested cells were sorted using MoFlo XDP (Beckman) and collected for RNA extraction.

### Immunofluorescence

For immunofluorescent analysis, kidneys were harvested and fixed in 4% formaldehyde overnight at 4°C. and then embedded in OCT to receive the frozen-section at 100 μm on the Micron HM550 cryostat. Primary antibodies used were anti-Cox2 (100034-lea, Cayman), β-catenin (51067-2-AP, Protein tech). The secondary antibodies were used at 1:500 (Goat Anti-Mouse IgG H&L Alexa Fluor® 647 (ab150115 Abcam)), (Donkey anti-Goat IgG (Alexa Fluor 647), (ab150131, Abcam)). The images were taken on the Nikon A1 confocal microscope.

### Statistics

All data are presented as means ± standard error of the mean (SEM). Except for special instructions, all experiments were carried out more than 3 independent experiments. Statistical analysis was performed using GraphPad Prism (version 8.02) and Excel (version, Microsoft office home and student 2019) for Microsoft Windows. To compare results from experiments involving two independent variables, we used two-way ANOVA, followed by the Bonferroni pairwise multiple comparison post-test.

## Acknowledgments

We thank Dr. Neil Hukriede for sending us the *lhx1a:EGFP*/pI-SceI plasmid. This work was funded by The National Key Research and Development Program of China (2017YFA0106600), the National Natural Science Foundation of China (No. 32070822, 31771609).

## Author contributions

C.L., T.Z., and J.Z. conceived the study; C.L., and X.L. designed the study; X.L., X.T., T.Y., D.J., Y.Z., S.L., and W.Y. performed all the experiment; X.L., C.L., and J.Z. carried out microscopy; C.L., and X.L. prepared the manuscript; all authors approved the final version.

## Data and materials availability

All data are available in the main text or the supplementary materials.

