## Supplementary Information for "Renal Interstitial Cells Promote Nephron Regeneration by Secreting Prostaglandin E2"

**This PDF file includes:**

Figure 1—figure supplement. Expression analysis of *col1a1b*; *col1a2*; *col6a4a*; *mmp2*.
Figure 2—figure supplement. RICs can be labeled by *Tg(fabp10a:GFP)* line.
Figure 3—figure supplement 1. Kidney injury promotes RICs to synthesize and secrete PGE2.
Figure 3—figure supplement 2. PGE2 receptors expression in *lhx1a*+ cell aggregates.
Figure 5—figure supplement 1. PGE2-EP4 signaling promote nephron regeneration through Wnt signaling.
Figure 6—figure supplement 1. RICs promote mesonephric development through synthesize and secret PGE2.
Legends for Figure 6—movie supplement 1
Legends for Figure 1—source data 1

**Other supplementary materials for this manuscript include the following:**

Figure 6—movie supplement 1. 3D movie of RICs wrapping renal progenitor cell aggregate corresponding to Figure 6F.
Figure 1—Source data 1. Single-cell RNA sequencing gene expression for each identified kidney cell population.
Supplementary Figures and Legends

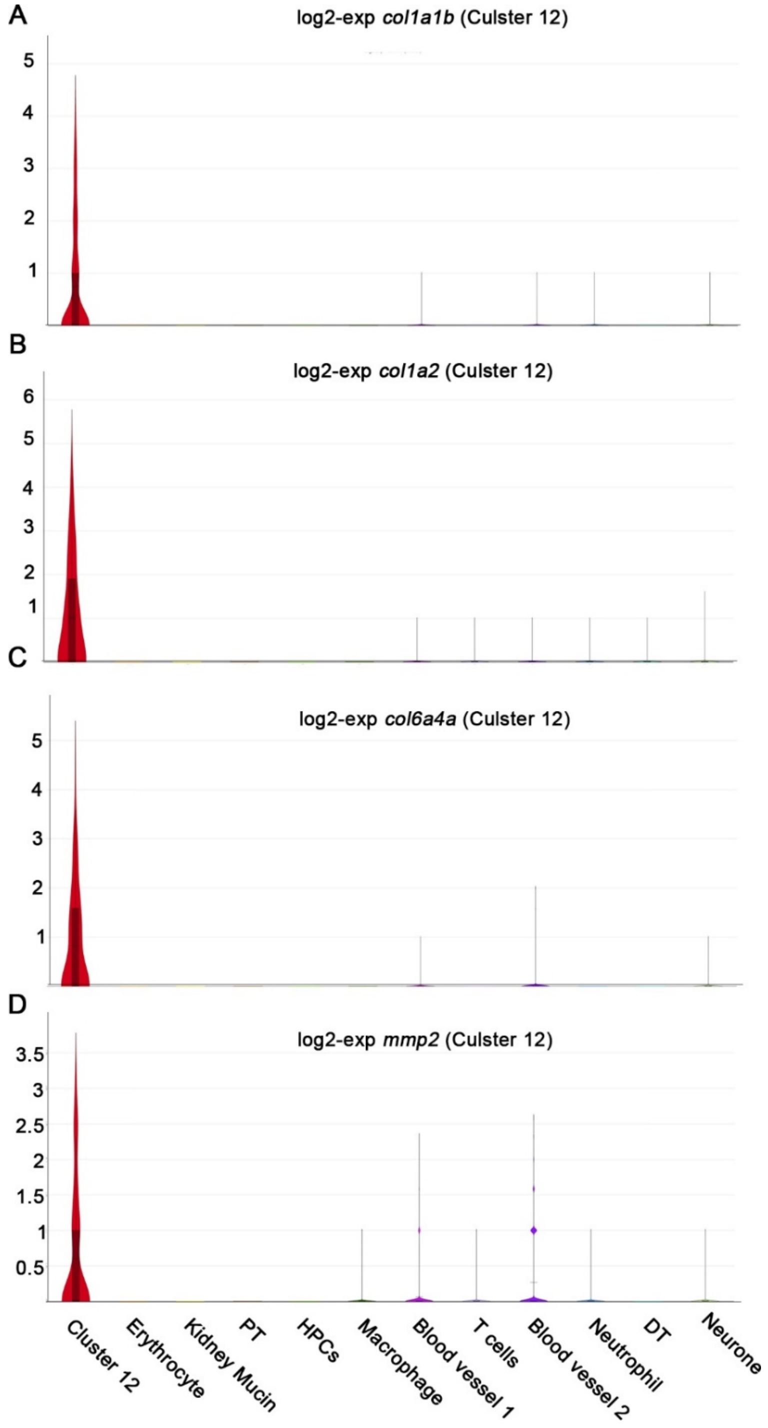


**Figure 1—figure supplement**. Expression analysis of *col1a1b* (**A**); *col1a2* (**B**); *col6a4a* (**C**); *mmp2* (**D**); Distal tubule; PT, Proximal tubule; HPCs, Hematopoietic stem cells.

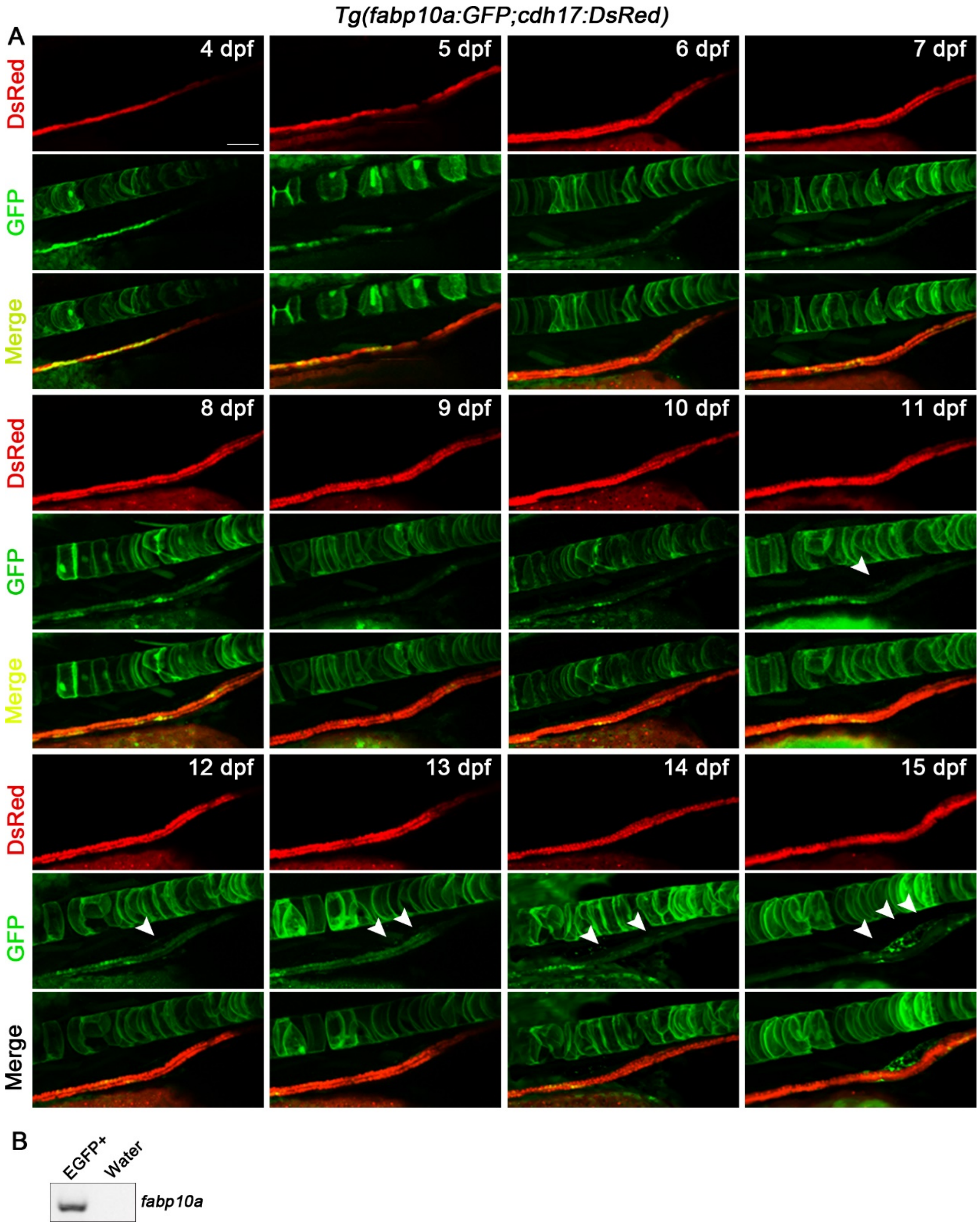


**Figure 2—figure supplement 1. RICs can be labeled by *Tg(fabp10a:GFP)* line.** (**A**) The development of *Tg(cdh17:DsRed; fabp10a:GFP)* line from 4 dpf to 15 dpf. *Tg(fabp10a:GFP)* label pronephric tubule in early stage and label RICs (arrowheads) after 11 dpf (n=5). (**B**) PCR analysis the expression of *fabp10a* in *Tg(fabp10a:GFP)* labeled kidney cells. *fabp10a* truly expressed in these cells. Scale bar in A, 100 μm.

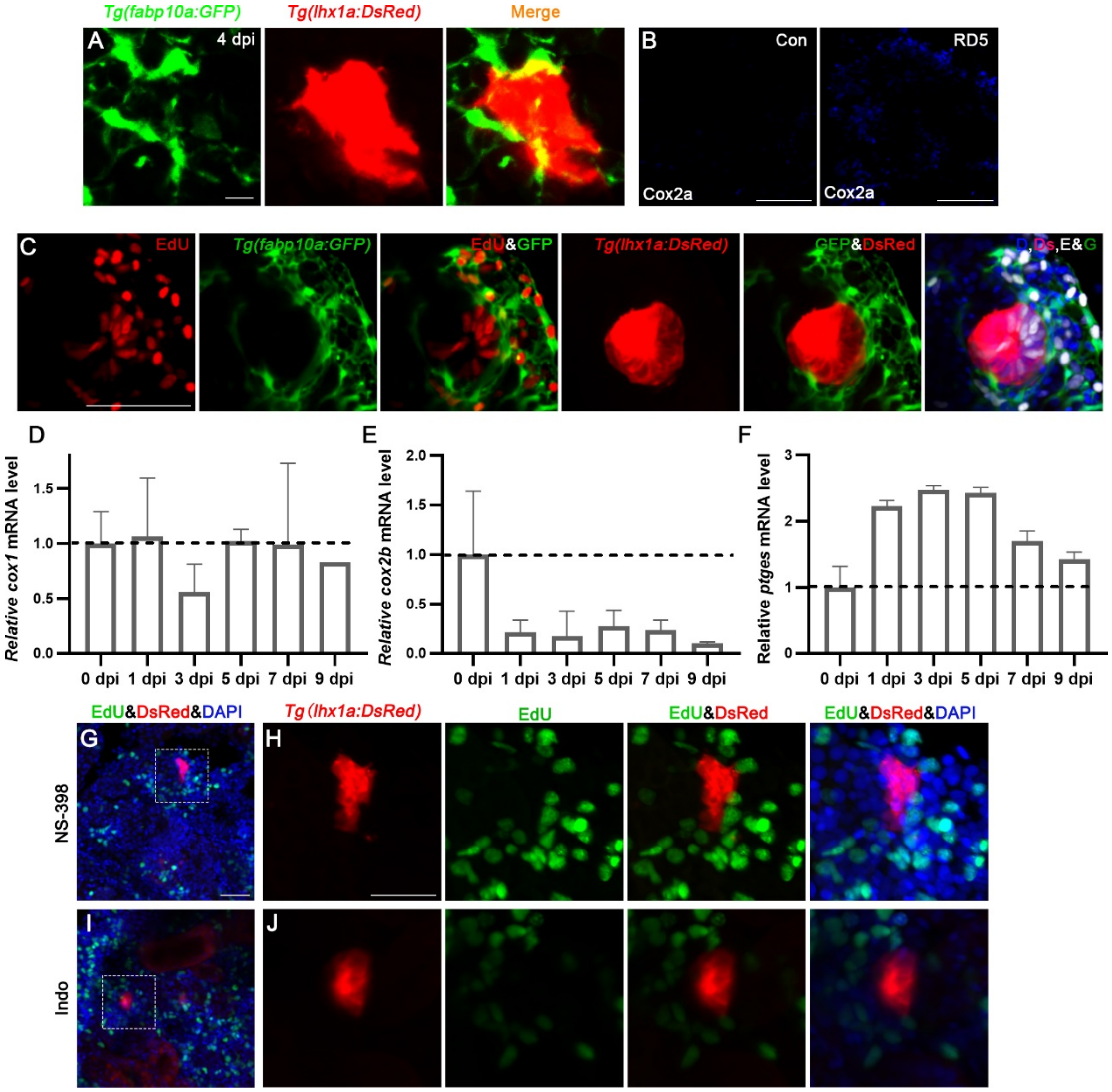


**Figure 3—figure supplement 1. Kidney injury promotes RICs to synthesize and secrete PGE2.** (**A**) Confocal stack projection of *Tg(fabp10a:GFP; lhx1a:DsRed)* transgenic kidney tissue at 4 dpi. RICs were wrapped around and closely interacted with the *lhx1a*$^{+}$ cell aggregate (n=4). (**B**) Proliferation assay of RICs which wrapped the *lhx1a*$^{+}$ cell aggregates at 5 dpi. D, DAPI; Ds, DsRed; E, EdU; G, GFP in merged image (n=6). (**C**) Immunofluorescent staining of Cox2a in kidney at 0 dpi (con) or 5 dpi. Cox2a was increased at 5 dpi (n=4). (**D-F**) qPCR relative quantification of *cox1a, cox2 and ptges* mRNA in kidney tissue harvested 0, 1, 3, 5, 7, 9 dpi. n=3 for each condition. (**G-J**) Intraperitoneal injection NS-398 or Indo reduces the proliferation of *lhx1a*$^{+}$ cell aggregates. H and J showed the higher-magnification image of boxed area shown in G and I. n=4 for each condition. Scale bar in A, 10 μm; B and C, 100 μm; G-J, 50 μm

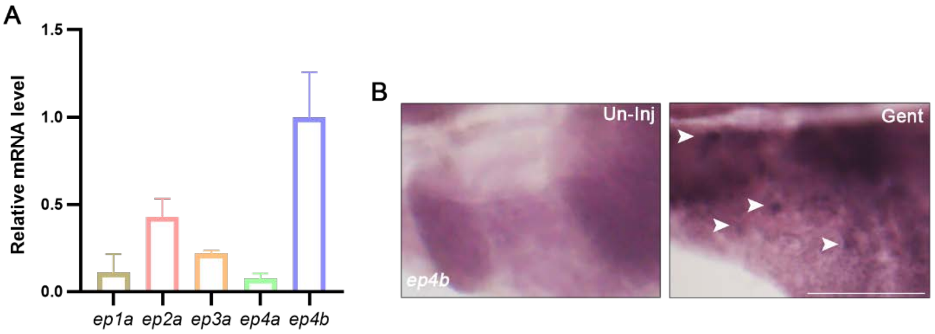


**Figure 3—figure supplement 2. PGE2 receptors expression in *lhx1a*⁺ cell aggregates.** (**A**) qPCR relative quantification of PGE2 receptors mRNA in *lhx1a*$^+$ cell. Results for each gene are normalized to *ep4b* and are presented as mean ± SEM of three independent experiments. (**B**) Whole-mount *ep4b in situ* hybridization showing the un-injured or injured zebrafish kidney region. *ep4b* labeled cell aggregates at 7 dpi. Scale bar in B, 500 µm.

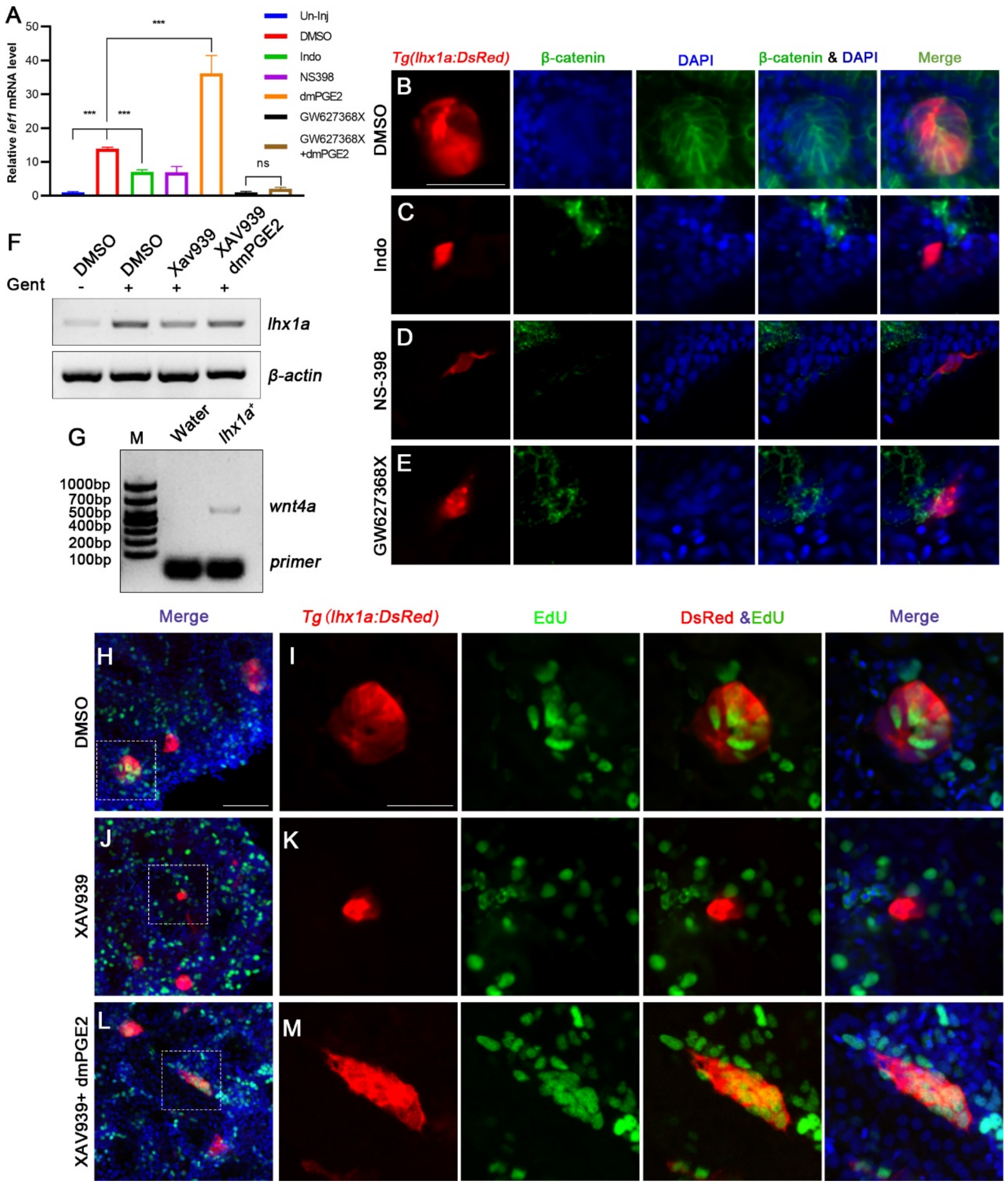


**Figure 5—figure supplement 1. PGE2-EP4 signaling promote nephron regeneration through Wnt signaling.** (**A**) qPCR relative quantification of *lef1* mRNA in kidney tissue harvested 7 dpi with different treatment. Data were analyzed by ANOVA, **$p < 0.01$, ***$p < 0.001$, ns, no significant difference (n=3 for each condition). (**B-E**) Immunofluorescent staining of β-catenin in *Tg(lhx1a:DsRed)* zebrafish kidney at 5 dpi. (**B**) Injected DSMO as control group and amount of β-catenin can be detected in *lhx1a*$^{+}$ cell aggregates cytoplasm and nucleus (arrowheads). Injected Indo (**C**), NS-398 (**D**) and GW627368X (E) can decrease β-catenin level in *lhx1a*$^{+}$ cell aggregates (n=4-6 in B, C, D and E). (**F**) The *lhx1a* mRNA level were determined by RT-PCR at 7 dpi in XAV939 or Both dmPGE2 and XAV939 treated kidney. (**G**) The *wnt4a* mRNA level were determined by RT-PCR in un-injured or injured kidney. *wnt4a* mRNA increased

at 7 dpi. (**H, I**) Gentamicin induces *lhx1a*$^{+}$ new nephrons with proliferating EdU$^{+}$ nuclei. (**J, K**) Intraperitoneal injection of XAV939 reduces the proliferation of *lhx1a*$^{+}$ cell aggregates. (**L, M**) dmPGE2 can rescue the inhibition of XAV939 and recovered the proliferation of *lhx1a*$^{+}$ cell aggregates (n=4 for each condition). I, K, and M showed the higher-magnification image of boxed area shown in H, J and L. Scale bar in H, J and L, 100 μm; I, K and M, 50 μm.

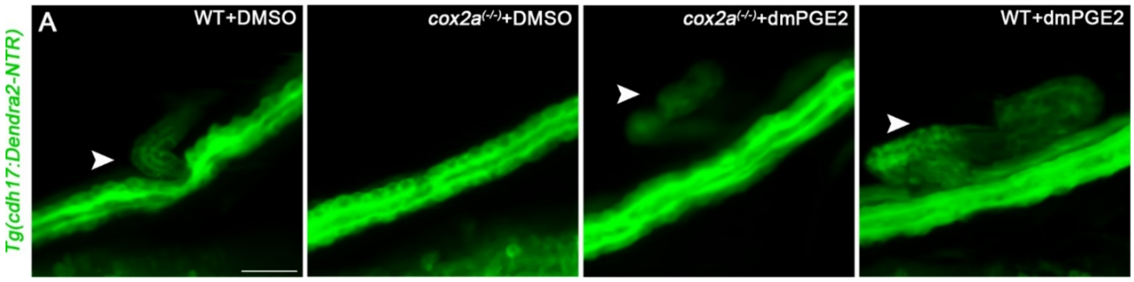


**Figure 6—figure supplement 1. RICs promote mesonephric development through synthesize and secret PGE2.** (**A**) *Tg(cdh17:Dendra2-NTR)* lines were treated with different condition at 4.6 mm stage and observed the mesonephric branch at 5.3 mm stage. The mesonephric development was inhibited in *cox2a* mutant and dmPGE2 can rescue the inhibition. The mesonephric development was promoted by exogenous dmPGE2 treatment. Scale bar, 100 μm (n=6).

**Figure 6—Movie supplement 1.** 3D movie of RICs wrapping renal progenitor cell aggregate corresponding to Figure 6F. Scale bar, 50 μm.

**Figure 1—source data 1.** Single-cell RNA sequencing gene expression for each identified kidney cell population.
